# Widespread inter-individual gene expression variability in *Arabidopsis thaliana*

**DOI:** 10.1101/335380

**Authors:** Sandra Cortijo, Zeynep Aydin, Sebastian Ahnert, James Locke

**Affiliations:** The Sainsbury Laboratory, University of Cambridge, Cambridge, CB2 1LR, UK

## Abstract

A fundamental question in biology is how gene expression is regulated to give rise to a phenotype. However, transcriptional variability is rarely considered and could influence the relationship between genotype and phenotype. It is known in unicellular organisms that gene expression is often noisy rather than uniform and this has been proposed to be beneficial when environmental conditions are unpredictable. However, little is known about transcriptional variability in plants. Using transcriptomic approaches, we analysed gene expression variability between individual *Arabidopsis thaliana* plants growing in identical conditions over a 24 hour time-course. We identified hundreds of genes that exhibit high inter-individual variability and found that many are involved in environmental responses. We also identified factors that might facilitate gene expression variability, such as gene length, the number of transcription factors regulating the genes and the chromatin environment. These results shed new light on the impact of transcriptional variability in gene expression regulation in plants.

## Introduction

Gene expression in individual cells is often noisy and dynamic. Genetically identical cells under the same environment can display widely different expression levels of key genes^1–3^. Noise in gene expression has been shown to have a significant impact on the design and function of genetic circuits in unicellular organisms^4,5^, and has been observed in multiple pathways in cell cultures of mammalian cells^6^. However, gene expression variability has only been analysed for a few individual genes in plants^7–9^. It is not known at a genome-wide scale to what extent gene expression can be variable during plant development.

Plants are a promising system to examine the global properties of noise in gene expression, as phenotypic variability, also referred to as phenotypic instability, has been observed in multiple areas of plant growth and development. This variability can occur both within as well as between individuals that are growing in the same conditions. High levels of phenotypic variability have been described for seed germination time^10–12^, patterning of lateral roots^13^ as well as for floral and foliar development^14,15^. Differences in the level of inter-individual variability has been observed between natural accessions, in recombinant inbred lines and also in mutants for many traits such as growth, hypocotyl length, leaf and flower number, plant height and plant defence metabolism^16–20^. This suggests that such variability can be controlled or buffered by genetic factors. However, the molecular mechanisms underlying such inter-individual phenotypic variability are still poorly understood.

In this work, we analyse gene expression variability between multicellular individuals using the model plant *Arabidopsis thaliana*, with the emphasis on three questions. Firstly, what is the global extent of gene expression variability between individuals? In order to better understand how gene expression variability is controlled and its role in plant physiological and developmental responses, we first need to identify genome-wide the genes that are highly variable between individuals. Secondly, does inter-individual expression variability change through the diurnal cycle? It is known that the diurnal cycle influences expression level of up to 36% of the transcriptome^21,22^. However, little is known about the impact of the diurnal cycle on gene expression variability. Thirdly, what factors can regulate this inter-individual expression variability?

Using single seedling RNA-seq, we identified hundreds of genes that are highly variable between individuals in Arabidopsis and show that the level of variability changes throughout the diurnal cycle. To ensure accessibility and reusability of our data, we created an interactive web-application, in which the inter-individual gene expression variability through a diurnal cycle can be observed for individual genes (https://jlgroup.shinyapps.io/aranoisy/). Moreover, we show that highly variable genes (HVGs) are enriched for environmentally responsive genes and characterised by a combination of specific genomic and epigenomic features. We have revealed both the level and potential mechanism behind gene expression variability between individuals in Arabidopsis, allowing understanding of a previously unexplored aspect of gene regulation during plant development.

## Results

### Widespread expression variability in Arabidopsis seedlings through the day and night

In order to measure transcriptional variability between individuals, we generated transcriptomes for single *Arabidopsis thaliana* seedlings at multiple time points over a full day/night cycle (Fig 1A). To minimise any variability caused by external factors, these seedlings originated from the same mother plant, germinated at the same time and were grown in the same plate under controlled conditions (Methods). To analyse how transcriptional variability is influenced by diurnal cycles, we harvested seedlings every 2 hours across a 24 hours period (Fig 1A). ZT2 to ZT12 corresponding to the time-points harvested during the day, and ZT14 to ZT24 to the time-points harvested during the night, ZT12 and ZT24 being respectively harvested just a few minutes before dusk and dawn. In total, 168 transcriptomes have been analysed, that is, of 14 individual seedlings for each of the 12 time points.

**Figure 1:**
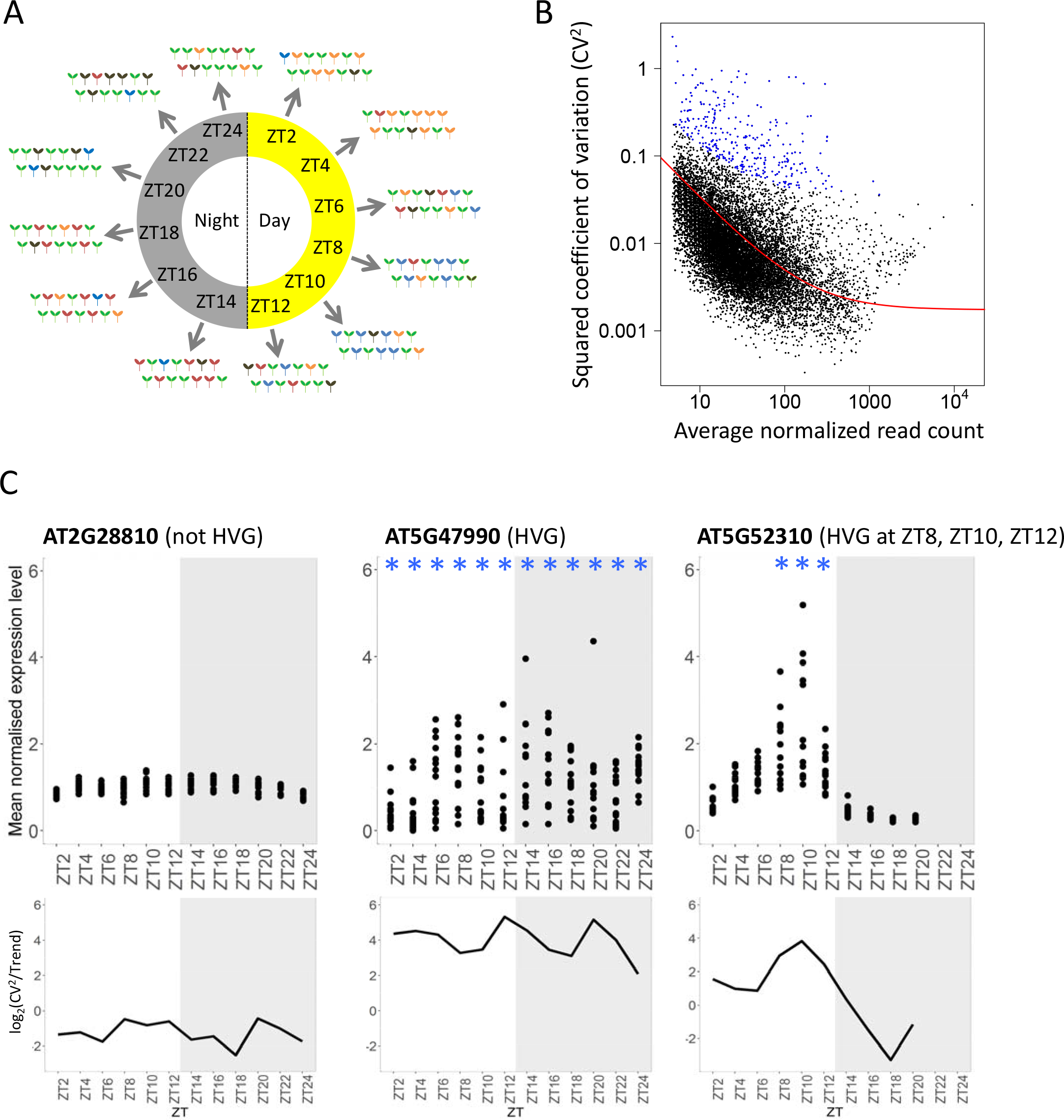
Widespread expression variability in *Arabidopsis seedlings* through the day and night. **A.** Experimental setup to identify transcriptional variability between seedlings during the day and night. RNA-seq was performed on individual seedlings, for a total of 14 seedlings at each time-point. Seedlings were harvested at 12 time-points, every 2 hours across a 24 hours period. Seedlings of different colors represent different transcriptional states, and thus inter-individual expression variability we aim to detect. **B.** Identification of highly variable genes (HVGs) for the time-point ZT2. The trend for the global relation (red line) between CV^2^ (variance/mean^2^) and mean expression is defined using all genes, minus small and lowly expressed genes (see material and method) and used to identify HVG (blue). For each gene, a corrected CV^2^ is calculated: log_2_(CV^2^ /trend). **C.** Expression profiles (top) in the 14 seedlings over a 24 hours time-course (with 12 time-points) for a non-variable gene (left), a highly variable gene (middle) and a gene with the level of variability changing across the 24 hours (right). Each dot is the mean normalised expression level for a single seedling. Variability profiles (bottom) of the log_2_(CV^2^ /trend) for the same genes are also shown. Blue stars indicate time-points for which the gene is detected as being variable in the RNA-seq.

For each time-point, we identified highly variable genes (HVGs) using a previously described method^23^ (Fig 1B). In order to avoid biases caused by technical noise, which is likely to be higher at lower expression levels, we only selected the HVGs if they were significantly more variable than the background trend in CV^2^ (Methods). We also calculated a corrected CV^2^ for each gene, log_2_(CV^2^/trend), which corrected for the observed negative trend between CV^2^ and expression level, and used it for further analyses of gene expression variability. Genes with a negative log_2_(CV^2^/trend) are less variable than the trend, while genes with a positive log_2_(CV^2^/trend) are more variable than the trend. We observed that while some genes are never classed as a highly variable gene (HVG) (AT2G28810, Fig 1C), others are selected as a HVG for the entire time course (AT5G47990, Fig 1C), or as a HVG for only some time points (AT5G52310, Fig 1C), indicating a broad range of variability profiles during the diurnal cycle.

We next verified that the variability detected using this approach could not be explained by experimental error or technical noise. First, in order to validate the profiles for gene expression variability during the time-course, we performed a full time-course replicate, and examined the variability between seedlings for 10 genes by RT-qPCR (Fig S1). These genes have been selected for not being highly variable (AT2G28810, At3G05880, AT1G30750, AT2G46830, AT4G27410, AT5G24470), being highly variable for the entire time-course (AT5G47990, AT5G15970), or highly variable for a few time-points of the time course (AT5G52310, AT1G08930, Fig S1). We observed very similar expression profiles for these genes (Fig S1A). We also found a positive correlation of 0.58 (Pearson test, p-value = 8.18e-11) between the CV^2^ measured by RNA-seq or RT-qPCR (Fig S1B), indicating that genes have a similar level of variability, relative to each other, in both experiments. We also obtained very similar CV^2^ at ZT24 when using an independent mapping programme (salmon instead of tophat, Fig S2B). To test if there were transcriptome-wide trends in the level of variability across the day, we verified that the global trends of the CV2 against the average normalised expression detected for each time-point are in the same range (Fig S2C). We also observed that there was no obvious bias in the distribution of CV^2^ against the average normalised expression at the different time-points (Fig S2D).

Having confirmed that the measured inter-individual variability is not due to technical noise or experimental error, we examined the HVGs in more detail. In total, we detected between 257 and 716 HVGs at each time point, with more HVGs detected during the night (Fig 2A). We also generated two other reference gene lists to compare to the HVGs: a randomly selected set of genes for each time-point, with the number of random genes selected matching the number of HVGs (Fig S3B), and the least 1000 variable genes for each time-point (LVGs, for lowly variable genes, see Fig S3A). We see that HVGs are at least 3 times and on average 9.3 times more variable than the global trend, while LVGs are at least 4.8 times and on average 8.9 times less variable than the global trend (Fig S2E). Random genes span a wide range of variability including values as low as for LVGs and as high as for HVGs. All these results taken together reveal a widespread level of inter-individual variability in gene expression throughout the entire time-course, with 8.7% of the analysed transcriptome detected as highly variable in at least one time-point. Moreover, as we describe in more detail below, some genes can show very different levels of variability during the time-course.

**Figure 2:**
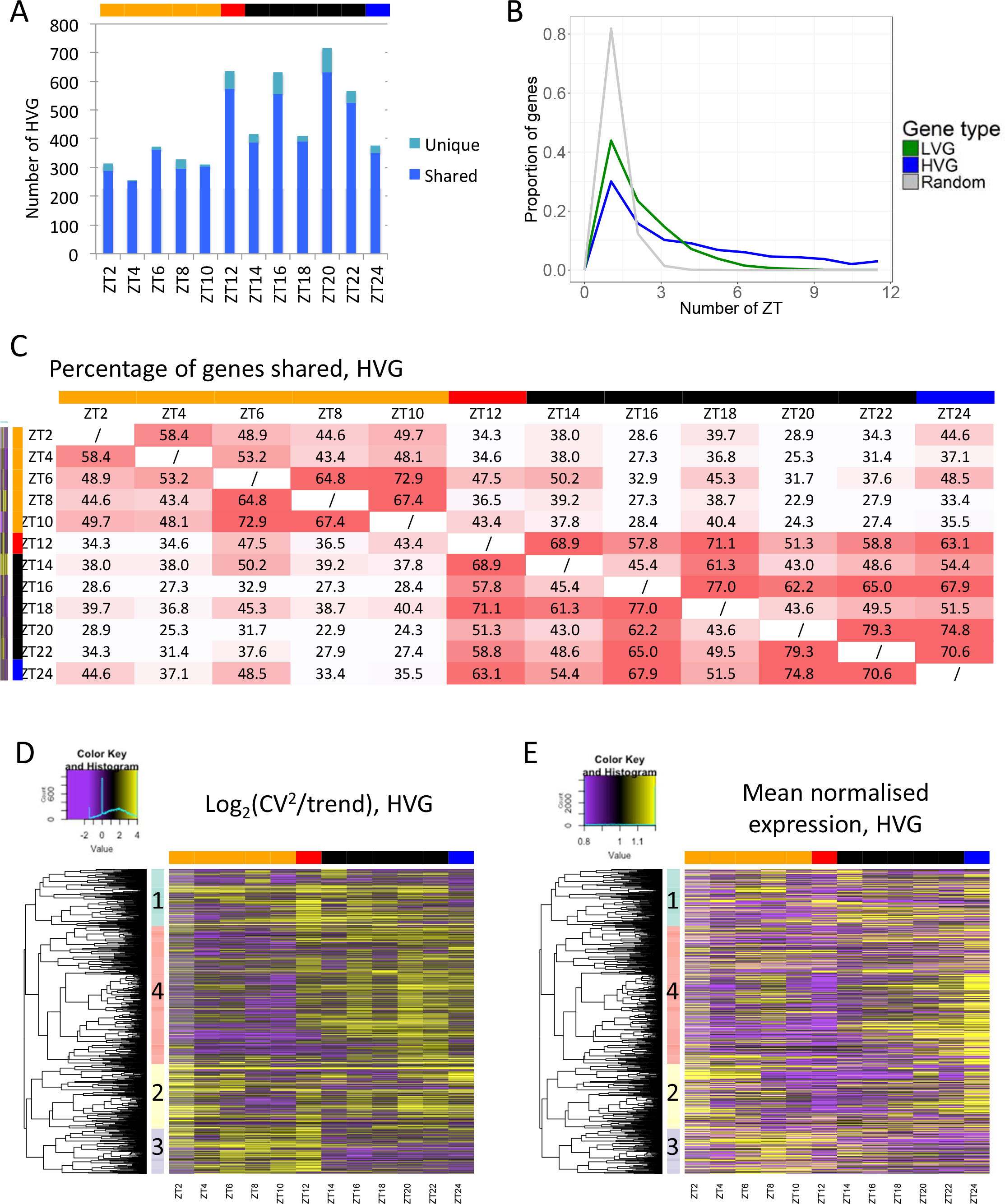
Structure of noisiness shows partitioning between day and night. **A.** Number of genes detected as being highly variable for each time-point. These genes are separated between those that are also selected as highly variable in at least one other time-point (dark blue) and those highly variable in only one time-point (light blue). The top bar indicates time-points harvested during the day (orange), just before dusk (red), during the night (black) and just before dawn (blue). **B.** Distribution of the number of time-points at which genes are identified as highly variable (blue), lowly variable (green) or are included in the random control set (grey). **C.** Heatmap of the percentage of highly variable genes shared between time-points. Red indicates a high percentage of HVGs in common between two time-points. The top and side bars indicates time-points harvested during the day (orange), just before dusk (red), during the night (black) and just before dawn (blue). **D.** Hierarchical clustering of HVGs based on the log_2_(CV^2^/trend) at each time point. The result is represented as a heatmap where yellow indicates a high log_2_(CV^2^/trend). The genes were separated into four clusters, indicated by the side colored bar. The top bar indicates time-points harvested during the day (orange), just before dusk (red), during the night (black) and just before dawn (blue). **E.** Heatmap of the normalised expression level for the genes in Fig 2D, keeping the same clustering organisation. The top bar indicates time-points harvested during the day (orange), just before dusk (red), during the night (black) and just before dawn (blue).

### Structure of noisy genes shows partitioning between day and night

We next examined the structure of our measured variability in more detail. First, we examined whether HVGs were scored as variable for multiple time-points, or for only one time-point. The vast majority (~93%) of the HVGs detected for each time-point are also detected as highly variable in another time-point (Fig 2A, dark blue). In comparison, an average of ~80% of LVGs and ~30% of random genes selected for each time-point are also observed in another time-point (Fig S3AB).

Since many genes are detected in more than one time point, we then defined in how many time-points they are identified. In total, 30% of all the 1358 HVGs are identified in only one time-point while the others are shared with other time-points, up to all of the 12 time-points for 40 genes (Fig 2B). LVGs and random genes are detected in a lower number of time-points, with 46% of all LVGs and 71% of all random genes that are specific for one time-point, while no genes are observed as lowly variable or random in all 12 time-points (Fig 2B). These results show that the number of HVGs shared between time-points is higher than what would be expected by chance and what is observed for LVGs. It indicates that genes could have profiles of variability across the time-course, as has been observed many times for mean gene expression levels, and be highly variable for multiple time-points.

Having observed that HVGs are shared between time-points, we then tested for any structure in the gene expression variability across the diurnal cycle. We calculated the percentage of HVGs that are shared between any two time-points and can see a clear separation of day and night time-points (Fig 2C). ZT12, which was harvested just a few minutes before dusk, behaves differently as it shares a high proportion of HVGs with night time points. When excluding ZT12, the percentage of HVGs that are shared between two time-points of the day (~55% on average), or two time-points of the night (~60%) is higher that between one time-point of the day and one time-point of the night (~35%). When doing the same analysis for LVGs, we observed that the percentage of genes that are shared between two time-points of the day (~18.5%), or two time-points of the night (~20.8%) is very similar to the percentage of genes shared between one time-point of the day and one time-point of the night (~17%, Fig S3C). We could not find any difference in the percentage of genes that are shared between two time-points in these three categories for random genes (Fig S3D). This result indicates a structure of the HVGs, but not of LVGs and random genes, in the time course, with a separation between day and night.

In order to identify profiles of inter-individual variability across the time course, we performed hierarchical clustering of all HVGs that were identified in at least one time-point (1358 genes) based on their log_2_(CV^2^/trend) at each time-point. We detected 4 clusters of variability patterns across the time course (Fig 2D). Two clusters (543 genes, clusters 1 and 2) are composed of genes being variable during the day and the night. One cluster (615 genes, cluster 4) is composed of genes being highly variable mainly during the night, while another one (200 genes, cluster 3) is composed of genes being highly variable mainly during the day. This observation is specific for HVGs, as we cannot observe such marked structure of variability profiles for LVGs and random genes (Fig S3E-F). All these results show a clear structure in gene expression variability between seedlings during the time course, with different sets of genes being variable during the day or during the night, further suggesting regulation of the level of variability.

### Highly variable genes are enriched in environmentally responsive genes

We next examined the function of the HVGs and LVGs. To do so, we first analysed Gene Ontology (GO) enrichment for all HVGs (1358 genes). We identified enrichment for several processes involved in the response to biotic and abiotic stresses as well as in the response to endogenous and exogenous signals (Table S1). This is not the case for the LVGs (5727 genes), for which we found enriched GOs involved in primary metabolism (Table S2), or for the random genes (4596 genes), for which no GO term was enriched.

Interestingly, different GOs are enriched in the clusters identified in Fig 2D based on the log_2_(CV^2^/trend) of HVGs along the time course (Table S1). For example, the response to cold is enriched in clusters 1 and 3, containing genes highly variable during the day, while nitrate assimilation is only enriched in cluster 4, which contains genes highly variable during the night (Table S1). This result suggests that some GOs might be variable at specific times of the diurnal cycle. In order to test this, we analysed GO enrichment for the HVGs identified at each time-point (Fig 3A), and clustered the GOs based on the log_10_(FDR) of their enrichment at the different time-points. While some GOs such as lipid transport and defence response to fungus are enriched in HVGs throughout the entire time course, we also identified GOs that are enriched only for a subset of the time course. This is the case for the response to toxic substance, reactive oxygen species metabolic process and response to iron ion that are more enriched during the night, or the response to water deprivation and to cold that are more enriched during the day (Fig 3A). We also analysed GO enrichment for the LVGs at each time-point and do not observe such enrichment of GOs preferentially during the day or night (Fig 3B, Table S2). We also observed that HVGs tend to be expressed with a higher tissue-specificity compared with LVGs and random genes (Fig S4), in agreement with the enrichment of many GOs associated with tissue-specific functions in HVGs.

**Figure 3:**
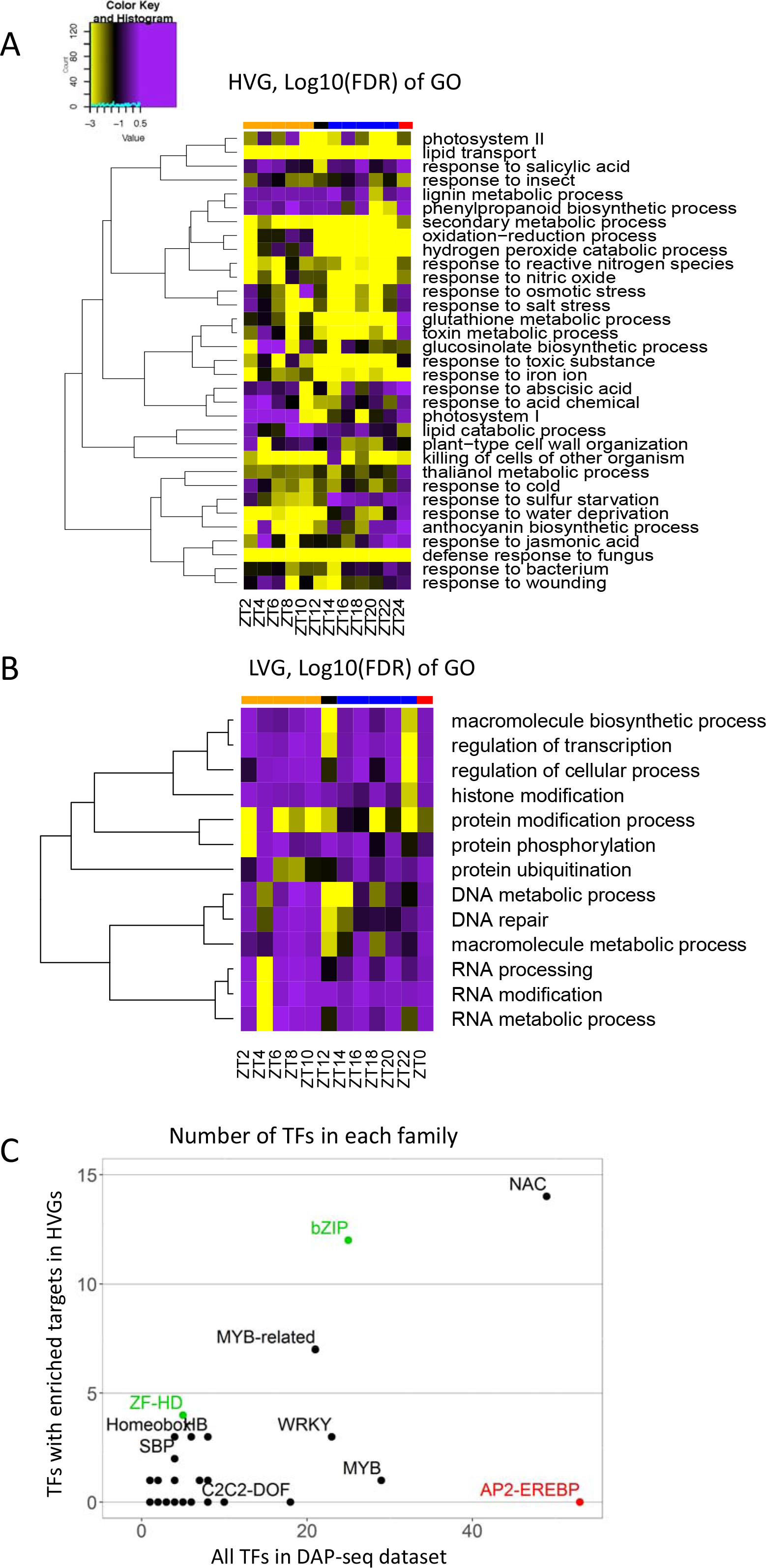
HVGs are enriched for stress responses. **A.** GO enrichment for genes selected as highly variable for each time-point. GOs that are enriched in at least one time-point are represented. Hierarchical clustering of the GO is performed on the log_10_(FDR) for GO enrichment in the HVGs. The result is presented as a heatmap with significantly enriched GO in yellow. **B.** GO enrichment for genes selected as lowly variable for each time-point. GOs that are enriched in at least one time-point are represented. Hierarchical clustering of the GO is performed on the log_10_(FDR) for GO enrichment in the LVGs. The result is presented as a heatmap with significantly enriched GO in yellow. **C.** Number of transcription factors (TF) in each TF family with enriched targets in the HVGs compared with the total number of TFs in each family included in the DAP-seq data. Families in green have a significantly higher number of TFs with enriched targets in the HVGs than in the entire dataset. Families in red have a significantly lower number of TFs with enriched targets in the HVGs than in the entire dataset.

In order to support these GO enrichment results, we also examined the transcription factors binding to the HVGs and LVGs, using available data generated by DNA affinity purification coupled with sequencing (DAP-seq), which provides the list of *in vitro* targets for 529 TFs^24^. We identified 60 TFs with enriched targets in the HVGs, 5 TFs with enriched targets in the LVGs and only one TF with enriched targets in the random genes. The 60 TFs with enriched targets in the HVGs are mainly part of the NAC, bZIP and MYB-related families. ZF-HD (4 TFs with enriched targets in the HVGs) and bZIP (12 TFs with enriched targets in the HVGs) families are significantly more represented in this set than expected based on the entire DAP-seq dataset (Fig 3C). bZIP TFs regulate multiple processes including pathogen defence, light and stress signalling, seed maturation and flower development^25^. These are in agreement with the enriched GOs identified for HVGs, involved in responses to the environment as well as biotic and abiotic stresses.

These results show that HVGs are enriched for genes involved in the response to environment and stress and are targeted by TF families involved in environmental responses, while LVGs are enriched in metabolism. Moreover, the clear pattern of enrichment for some function either during the day or the night further suggests that variability between seedlings across the day and night is functional and might be controlled.

### Gene expression profiles and variability are not correlated for the majority of the HVGs

Having identified that HVGs are enriched in stress responsive genes and that variability is structured during a diurnal cycle, we next asked, what factors could be involved in modulating variability in expression? To test if expression levels could modulate variability, we analysed the expression level of HVGs, LVGs and random genes at each time-point (Fig S5A). We observed that HVGs and random genes have very similar expression levels, as expected by the fact that we took into account the negative trend between CV^2^ and expression when selecting HVGs. Their expression levels are also similar but slightly higher than LVGs. In order to define if expression profiles could influence changes in variability during the time course, we used the same hierarchical clustering of HVGs based on their log_2_(CV^2^/trend) (Fig 2D) to represent their mean normalised expression levels (Fig 2E). We observed that genes in cluster 4, which are more variable during the night, have a peak of expression at the end of the night. Genes in cluster 3, that are more variable during the day, are also slightly more expressed during the day. However, at a global scale we cannot see a general link between profiles of gene expression level and variability for all HVGs (Fig 2E). To go further, we analysed the correlation between the profiles of mean normalised expression and of the log_2_(CV^2^/trend) profile for each of the 1358 HVGs, 5727 LVGs as well as for a set of 1358 random genes (Fig S4B). We observed a slightly higher correlation for HVGs (median of 0.19) compared to LVGs (median of -0.06) and random genes (median of 0.009). However, while variability and expression levels are positively correlated for some HVGs (peak around 0.5 in Fig S5B) this is not the case for many other HVGs (peak around 0 in Fig S5B). If we consider both positive (> 0.4, example Fig S5C) and negative (< -0.4, example Fig S5D) correlation, it seems that profiles in gene expression variability for approximately 45% of HVGs could be potentially explained by changes in expression levels during the time-course. No clear link can be observed between gene expression and variability profiles for the remaining 55% of the genes (example Fig S5E). Altogether, these results suggest that profiles in variability could potentially be explained by changes in expression level for only half of HVGs, indicating that other factors might be involved in facilitating gene expression variability.

### Noisy genes tend to be smaller and to be targeted by more Transcription Factors

In order to identify other factors that might be involved in regulating gene expression variability, we analysed several genomic features including gene length, number of introns and the number of TFs targeting the genes. We first observed that HVGs tend to be shorter and contain a lower number of introns than LVGs or random genes (Fig 4A-B, Fig S6A-B). We also observed a negative correlation between the level of variability and the gene length or number of introns for all genes at each time-point (Fig S6C-D). As the gene length and number of introns are strongly positively correlated (Fig S6E), we analysed the impact of one of these factors on gene expression variability while fixing the other and vice-versa. We observed very similar distributions for the number of introns of HVGs, LVGs and random genes when these genes are of similar size (Fig S6F). On the contrary, we still observe a trend for HVGs to be smaller when comparing genes with the same number of introns, for genes with 3 introns and less (Fig S6G). These results suggest that gene length might have a more important role than the number of introns in facilitating gene expression variability. In order to check for potential experimental bias that could account for the fact that smaller genes are more variable, we fragmented *in silico* 27 genes of ~1.5 to ~2.5kb into smaller fragments of ~250-300bp and examined if this could affect the level of gene expression variability that we estimate (Fig S6H). We performed this analysis for genes with different levels of expression that are either HVGs, LVGs or have a corrected CV^2^ around zero (i.e. close to the global trend). We observed a very similar level of corrected CV^2^ for full genes and their fragments (Fig S6H). Only 2 fragments out of the 35 (5%) originating from HVGs are not any more detected as highly variable, and only 2 fragments out of the 63 (3%) originating from genes that are not highly variable are now detected as highly variable. These results suggest that the trend we observe of HVGs to be smaller is not caused by technical biases.

**Figure 4:**
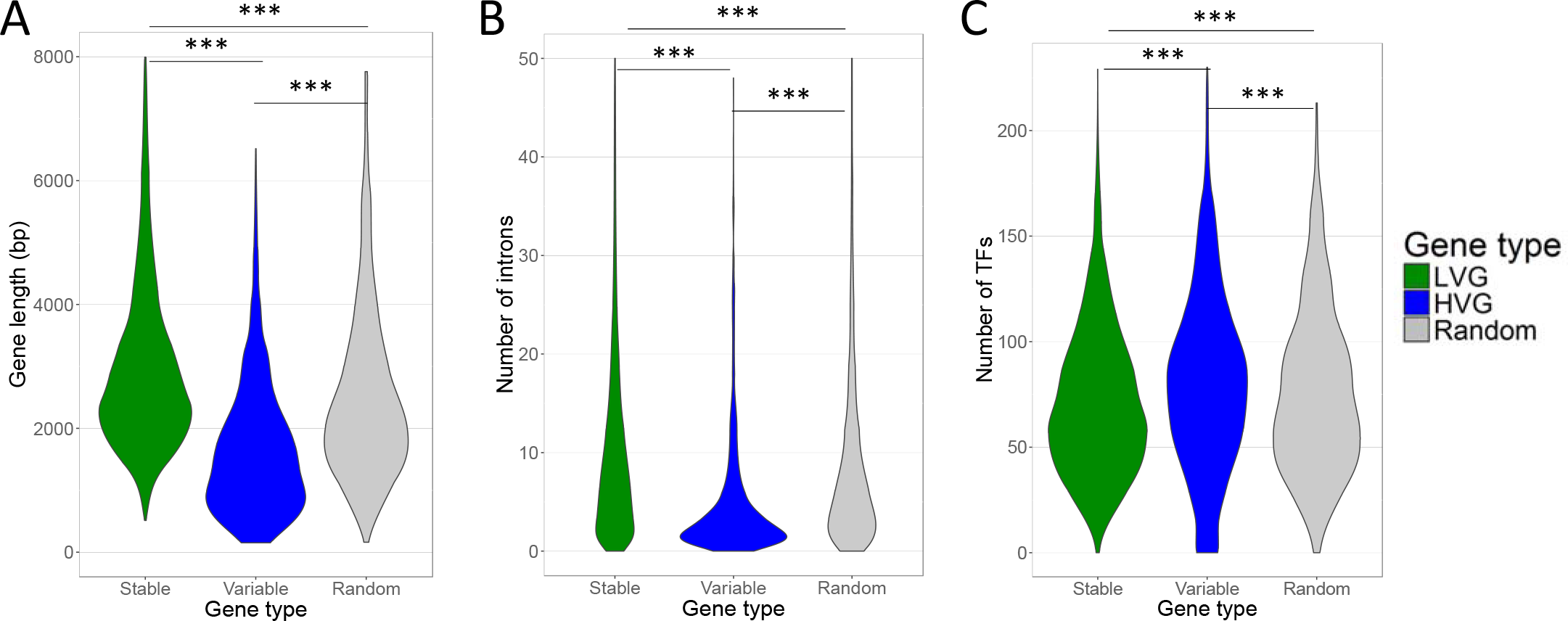
HVGs tend to be small and to be targeted by a higher number of TFs. **A.** Distribution of the gene size for LVGs (green), HVGs (blue) and other genes (grey). *** Indicates that gene size is significantly different with a p-value < 0.001. **B.** Distribution of the number of introns for LVG (green), HVG (blue) and other genes (grey). *** Indicates that the number of introns is significantly different with a p-value < 0.001. **C.** Distribution of the number of TFs targeting a gene, based on the DAP-seq available dataset, for LVG (green), HVG (blue) and other genes (grey). *** Indicates that gene size is significantly different with a p-value < 0.001.

One other factor we tested is the binding of transcription factors (TFs) at the promoters of genes. For this, we counted the number of TFs binding to the promoter of HVGs, LVGs and random genes using the available DAP-seq data and found a tendency for a higher number of TFs binding the promoter of HVGs (Fig 4C). This result suggests differences in the way HVG and LVG expression are regulated, which could possibly be due to different network architectures.

### Noisy genes tend to have a chromatin environment refractory to expression

On top of genomic features, another factor that can influence gene expression is the chromatin structure. In order to identify if HVGs are characterised by a specific chromatin structure, we analysed several histone marks using data already available. We first analysed the proportion of genes containing a histone modification among HVGs, LVGs and random genes in comparison with all background genes. We could identify that HVGs are enriched in H3K27me1, H3K27me3, which are repressive marks, while they are depleted in active marks such as H3K4me2, H3K4me3, H3K36me3 or H2Bub (Fig 5A).

**Figure 5:**
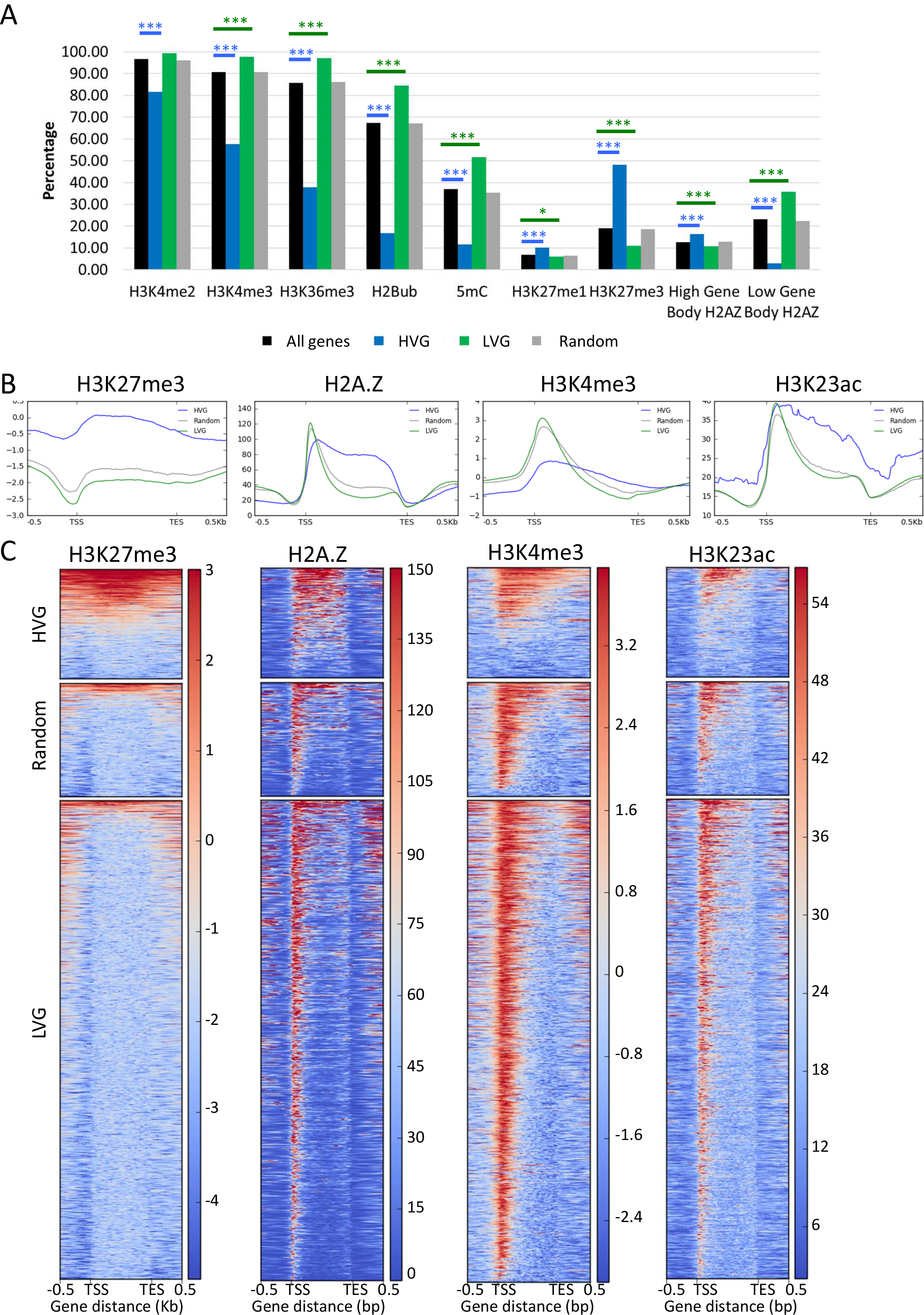
HVGs tend to have a specific chromatin environment. **A.** Proportion of genes marked with several chromatin marks among all genes passing size and expression level thresholds (black), HVGs (blue), LVGs (green) and random genes (grey). Blue and green stars indicate statistical differences in the number of marked genes compare to all genes, for variable and stable genes respectively (* Indicates a p-value < 0.05, *** Indicates a p-value < 0.001). **B.** Average profile for H3K27me3, H2A.Z, H3K4me3 and H3K23ac at HVGs (blue), LVGs (green) and random genes (grey). **C.** Heatmap of the enrichment for H3K27me3, H2A.Z, H3K4me3 and H3K23ac for HVGs (top), random genes (middle) and LVG (bottom). Red means a high level and blue means a low level for the chromatin marks.

They are also depleted in DNA methylation (Fig 5A), which is usually considered as a permissive mark for expression when in the body of genes^26,27^. On the other hand, LVGs are enriched in these active marks and depleted in H3K27me1 and H3K27me3. From previous studies, genes containing H2A.Z histone variants have been separated into two classes: (1) genes with a high signal in the gene body, which are enriched for environmentally-responsive genes and genes with tissue-specificity expression, and (2) genes with a low signal in the gene body for which H2A.Z is mainly observed at the 1st nucleosome, which are enriched for housekeeping genes^27^. The former category is enriched among HVGs, while the latter category is enriched among LVGs.

To define if HVGs and LVGs are also characterised by different profiles for these chromatin marks, rather than just differing in their presence/absence, we used already published ChIP-seq data for several chromatin marks and represented the signal along the genes. We identified differences in the profiles of the average chromatin signal between HVGs, LVGs and random genes for H3K27me3, H2A.Z, H3K4me3 and H3K23ac (Fig 5B). A higher H3K27me3 average signal is observed for HVGs (Fig 5B) and can be explained by a higher number of HVGs containing this mark compare to LVGs and random genes (Fig 5C). We observed H2A.Z and H3K23ac signal throughout the gene body for HVGs, while LVGs and random genes are characterised by a peak around the TSS, corresponding to the 1st nucleosome, and a lower signal for the rest of the gene body (Fig 5B-C). We see a higher H3K4me3 average signal for LVGs and random genes characterised with a peak at the beginning of the genes, while less than half of the HVGs have a high signal for these chromatin marks. To correct for differences in gene size between HVGs and LVGs, we also performed the same analysis on a subset of 150 HVGs and 185 LVGs and 125 random genes that have a similar size of 1100 to 1400 bp (Fig S7). The results are broadly the same as the ones obtained on all HVGs and LVGs.

These results indicate that HVGs and LVGs are characterised by a specific chromatin environment, in terms of the presence/absence of chromatin marks as well as for the profiles of these marks. They indicate that chromatin at HVGs tend to be more compacted and refractory to expression than at LVGs and random genes, which might have implications for how gene expression is regulated in these genes.

## Discussion

In this work, we have characterised the variability in gene expression between individual Arabidopsis seedlings at the genome-wide scale throughout a diurnal cycle. To do this, we have analysed 14 seedlings at each of the 12 time-points, generating 168 transcriptomes in total. This resource reveals previously unexplored variability for multiple pathways of interest for plant researchers, as well as providing insights into the modulation of gene expression variability at the genome-wide scale (Fig 6). We have successfully identified highly variable genes across the diurnal cycle, finding two sets of genes variable either during the day or night (Fig 2 and S3), revealed the functional classes of highly variable genes (Fig 3) as well as their genomic and epigenomic characteristics (Fig 4-5 and S6-S7). Interestingly, most profiles of gene expression variability are not correlated with profiles in gene expression levels (Fig S5), indicating that changes in expression levels during the diurnal cycle are not sufficient to explain changes in inter-individual variability. The large degree of gene expression variability revealed by our study will impact on our functional understanding of pathways as well as experimental design. To enable researchers to access this resource, we have created a graphical web interface to allow easy visualisation of inter-individual gene expression variability during a diurnal cycle for genes of interest (https://jlgroup.shinyapps.io/aranoisy/). This data could also be used for other purposes, such as inferring regulatory networks based on gene expression correlation between seedlings, as previously done using microarrays of individual leaves^28^.

**Figure 6:**
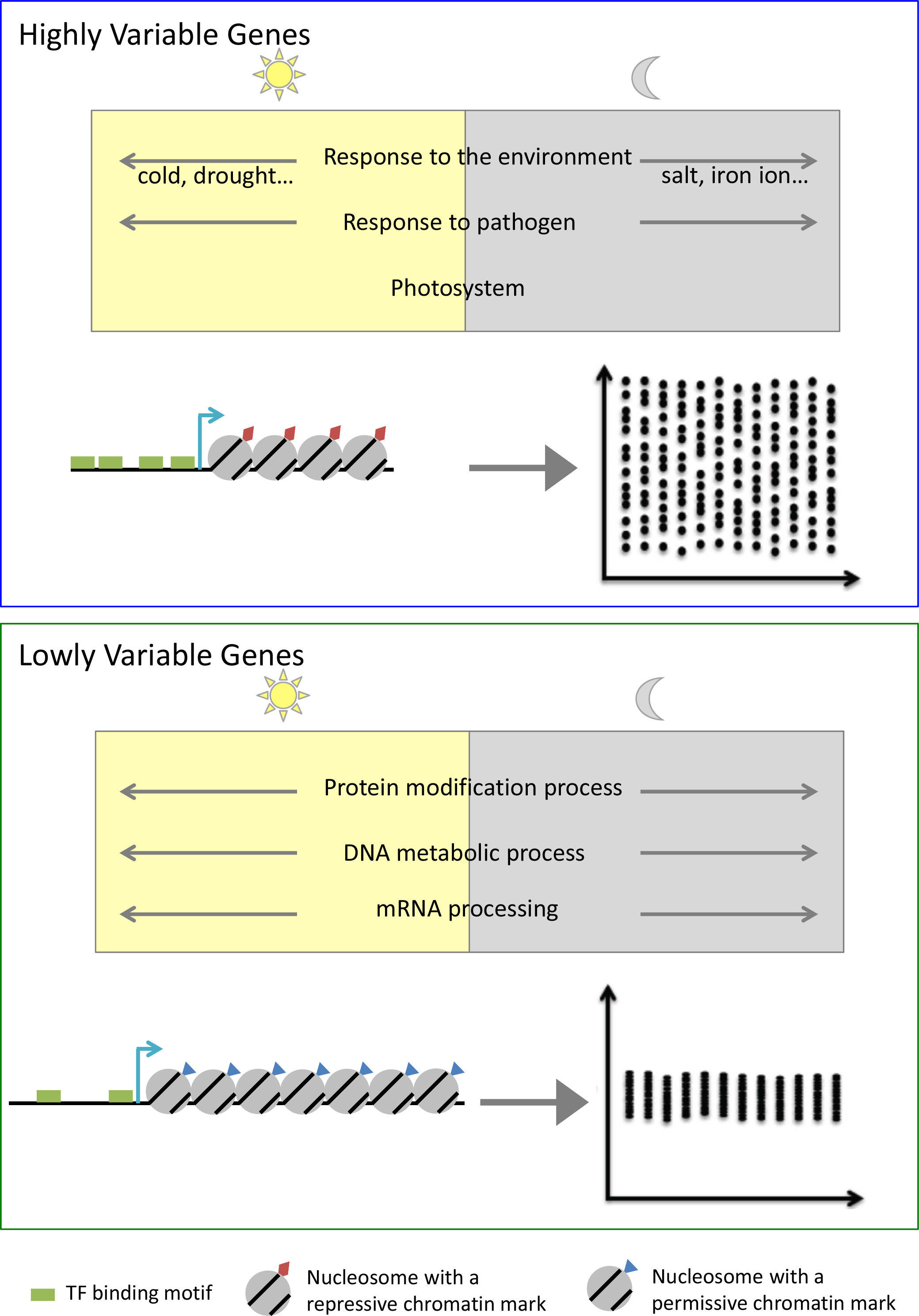
Models of HVGs and LVGs. HVGs (top panel) are enriched in environmentally responsive genes, and some functions are more variable either mainly during the day (yellow rectangle) or the night (grey rectangle). These genes tend to be smaller, to be targeted by a higher number of TFs (represented by the TFs binding motifs in green) and to be enriched in repressive chromatin marks (red rectangle on nucleosomes). On the other hand, LVGs (top panel) are enriched in genes involved in primary metabolism. These genes tend to be longer, to be targeted by a smaller number of TFs (represented by the TFs binding motifs in green) and to be enriched in permissive chromatin marks (blue triangle on nucleosomes), compared to HVGs.

We found that HVGs tend to be enriched for GOs involved in the response to environment, such as photosystems I and II, response to pathogens, response to abiotic stresses and response to iron ion. We also observed a high number of stress responsive TFs with targets enriched in HVGs. This is in agreement with previous observations in mammals and Yeast that HVGs are enriched in stress responsive genes^6,29,30^, and that LVGs are enriched in housekeeping genes^31^. This is also further supported by previous results showing a positive correlation between gene expression variability and plasticity^32^, the latter corresponding to environmentally triggered gene expression changes. It was also proposed in single celled organisms that transcriptional noise could be beneficial under unpredictable conditions^33–38^, a concept also known as bet hedging. In particular, gene expression variability for stress responsive genes between cells in a population was associated with survival of a fraction of cells during stress treatment and reconstitution of the full population once favorable conditions returned^39,40^. It is interesting to note that we have found functional classes of highly variable genes that are similar to the ones found for variable genes in single celled organisms. This is the case even though our work is at the whole plant scale, averaged over 10000s of cells, which suggests similar but different mechanisms for the generation of this transcriptional variability. Given the high number of environmentally responsive genes among HVGs, it would be of interest to test if inter-individual variability in stress responsive genes could be correlated with variation in stress survival in *Arabidopsis thaliana*. This hypothesis is probable, as phenotypic variability has been observed for many traits in *Arabidopsis thaliana*^13–17^. Moreover, the proportion of wild-type plants surviving to a stress is not zero in many studies^41–44^, suggesting the possibility of underlying gene expression variability for stress responsive genes explaining this observation. Our analysis of inter-individual gene expression variability was performed under non-stressed controlled conditions, but in the future it would be interesting to investigate how gene expression variability is influenced by changes in the environment and stress. Indeed, it was shown in yeast that genes coding for ribosomal proteins display a low level of variability in absence of stress but become more variable during stress treatment^29^. On the other hand, genes involved in environmental stress response are highly variable in the absence of stress but show a reduction of their variability during stress treatment^29^.

We identified several genomic and epigenomic factors that are correlated with gene expression variability. We found that HVGs tend to be shorter and targeted by a higher number of TFs than LVGs. In line with our results, a negative correlation was also previously observed in Yeast between gene length and noise for genes with a low plasticity^45^. It has also been shown in *Arabidopsis thaliana* that stress responsive genes are shorter^46^, in agreement with the fact that HVGs are enriched in environmentally responsive genes. Our results are further supported by previous studies showing that genetic factors can control or buffer inter-individual phenotypic variability in *Arabidopsis thaliana*^16–20^ Cis factors have also been shown to regulate gene expression variability in other organisms: TATA boxes are linked to gene expression variability in Yeast^47^, and the strength of cis-regulatory elements affects transcriptional noise in mammals^48^. The diurnal profiles we observed in inter-individual variability however indicate that genetic factors can only make genes prone to be variable but are not sufficient to explain their variability level at a given time of the day and that other factors, such as gene regulatory networks for example, are involved in modulating the level of variability of a gene.

On top of genomic factors, we identified that HVGs and LVGs have distinct chromatin profiles, with HVGs being characterised by an enrichment in H3K27me1 and H3K27me3, which are repressive marks, and depleted in active marks such as H3K4me2, H3K4me3, H3K36me3, H2Bub or DNA methylation. Chromatin has been shown to regulate the level of transcriptional noise, sometimes independently of expression level, in mammals^31,49^ and yeast^50^. In plants, over-expression of CHR23, a chromatin remodeler, is associated with an increase in inter-individual phenotypic and transcriptional variability^18^. These previous observations are in agreement with our results and suggest a role of the chromatin structure in regulating the level of gene expression variability, potentially with more compacted chromatin environments being more favorable to high variability. We nonetheless have to keep in mind that all these genomic and epigenomic factors are linked, as environmentally responsive genes have been shown to be smaller and to have a high gene body H2A.Z signal^27^, and that H3K27me3 was shown to be more enriched at small genes^51^. Further work perturbing these factors will thus be needed in order to decipher which ones are the major factors influencing inter-individual transcriptional variability.

## Material and method

### Plant materials and growth conditions

Col-0 WT *Arabidopsis thaliana* seeds were sterilised, stratified for 3 days at 4°C in dark and transferred for germination on solid 1X Murashige and Skoog (MS) media at 22°C in long days for 24 hours. Using a binocular microscope, seeds that were at the same stage of germination were transferred into a new plate containing solid 1X MS media. In total, 16 seeds were transferred into each of the 12 individual plates. Seedlings were grown at 22°C, 65% humidity, with 12 hours of light (170 μmoles) and 12 hours of dark in a conviron reach in cabinet. After 7 days of growth, seedlings were harvested individually into a 96-well plate and flash-frozen in dry ice. Sixteen seedlings were harvested at each time point, every two hours over a 24 hours period. In order to reduce environment effects, all seedlings harvested for one time-point were growing in the same plate, and seedlings that looked smaller than others were not harvested. Moreover, the seedling number corresponds to the seedling position in the plate and we could not see any obvious position effect when analysing gene expression variability. ZT2 to ZT12 corresponding to time-points harvested during the day, and ZT14 to ZT24 to time-points harvested during the night, ZT12 and ZT24 being respectively harvested just a few minutes after dusk and dawn (Fig 1A). Night time-points were harvested in the dark using a green lamp in order to avoid any interruption of the dark period with white light.

### RNA-seq library preparation

Sixteen seven-day old Col-0 WT Arabidopsis seedlings were harvested individually and flash-frozen in dry ice every two hours over a 24 hours period. Total RNA was isolated from 1 ground seedling using the MagMAX™-96 Total RNA Isolation Kit following manufacturer’s recommendation. RNA quality and integrity were assessed on the Agilent 2200 TapeStation, and RNA concentration was assessed using Qubit RNA HS assay kit. Library preparation was performed using the TruSeq Stranded mRNA library preparation kit (Illumina, RS-122-2101), for 1 μg of high integrity total RNA (RIN>8) into which 2 ul of diluted 1:100e ERCC RNA Spike-In Mix (ThermoFisher, cat 4456740) was added. The libraries were sequenced on a NextSeq500 using paired-end sequencing of 75 bp in length.

### RNA-seq mapping, detection of HVG and corrected CV2 calculation

The raw reads were analysed using a combination of publicly available software and in house scripts. We first assessed the quality of reads using FastQC (www.bioinformatics.babraham.ac.uk/projects/fastqc/). Potential adaptor contamination and low quality trailing sequences were removed using Trimmomatic^52^, before aligned to the TAIR10 transcriptome using Tophat^53^. Potential optical duplicates resulted from library preparation were removed using the Picard tools (https://github.com/broadinstitute/picard). For each gene, raw reads and TPM (Transcripts Per Million)^54^ were computed. Identification of HVGs was performed separately for each time-point as described previously^23^, using the code from M3Drop R package (https://github.com/tallulandrews/M3Drop, see detail in supplementary information). For each gene a corrected CV^2^ was calculated in order to correct for the negative trend observed between CV^2^ and the averaged expression level, as: log2(CV2/Trend for the same expression level)

### RT-qPCR

Sixteen seven-day old Col-0 WT *Arabidopsis thaliana* seedlings were harvested individually and flash-frozen in dry ice every two hours over a 24 hours period. Total RNA was isolated from 1 ground seedling. RNA concentration was assessed using Qubit RNA HS assay kit. cDNA synthesis was performed on 700ng of DNAse treated RNA using the Transcriptor First Strand cDNA Synthesis Kit. For RT-qPCR analysis, 0.4 μl of cDNA was used as template in a 10 μl reaction performed in the LightCycler 480 instrument using LC480 SYBR green I master. Gene expression relative to two control genes (SandF and PP2A) was measured. Then, in order to directly compare RT-qPCR data with RNA-seq data, expression levels for each gene were normalised by the averaged expression level of that gene across all seedlings at all time-points.

### Hierarchical clustering and principal component analysis

First a binary table was created for HVGs, LVGs and random genes indicating if a gene is selected (1) or not selected (0) as HVG, LVG or a random gene at each time-point. Using these tables, hierarchical clustering was performed using the statistical programme R (R Core Team 2015) using the function hclust on 1-pearson correlation. Principal component analysis (PCA) was performed in the statistical programme R (R Core Team 2015), using the function prcomp on the transposed binary table.

### Gene Ontology enrichment analysis

To assess over-represented biological functions of the genes in different clusters, we performed the GO enrichment analysis using the Gene Ontology enrichment analysis. GO enrichment p-values were calculated using Gene Ontology Consortium enrichment analysis tool (ref). All the genes for which a corrected CV^2^ was calculated were used as a background list, and this for each time-point separately, for the enrichment analyses.

### ChIP-seq mapping and profiling

The lists of genes being marked by the analysed chromatin marks were obtained from Roudier and colleagues^51^ and from Coleman-Derr and Zilberman^27^.

ChIP-seq data were downloaded from GSE101220 for H3K27me3^55^, from GSE79355 for H2A.Z^56^, from GSE73972 for H3K4me3^57^, and from GSE51304 for H3K23ac and H3^58^. Sequenced ChIP-seq data were analysed in house, following the same quality control and pre-processing as in RNA-seq. The adaptor-trimmed reads were mapped to the TAIR10 reference genome using Bowtie2^59^. Potential optical duplicates were removed using Picard, as described earlier. Averaged profiles and heatmap of the ChIP-seq signal along the gene body from 500bp upstream to the Transcription Start Site (TSS) to 500bp downstream of the Transcription Termination Site (TTS) were generated using deepTools^60,61^. The ChIP signal was normalised by the INPUT, when available (for H3K27me3 and H3K4me3).

## Data and code availability

Raw data are deposited on GEO (GSE115583)

Graphical web interface: https://jlgroup.shinyapps.io/aranoisy/

Codes used to analyse RNA-seq and ChIP-seq data are available upon request.

## Acknowledgements

We thank Dr Hugo Tavares (Sainsbury Laboratory Cambridge University, UK) for helping with the creation of the interactive web interface. We thank Dr Katie Abley (Sainsbury Laboratory Cambridge University, UK) and Dr Varodom Charoensawan (Mahidol University, Thailand) for critical reading of the manuscript draft. This research was made possible by a fellowship from the Gatsby Foundation (GAT3272/GLC). The work in the Locke laboratory is further supported by the European Research Council under the European Union’s Seventh Framework Programme (FP/2007-2013)/ERC Grant Agreement 338060.

## Author contributions

S.C. conceived of the project. S.C. and J.C.W.L. designed the project. S.C. performed RNA-Seq experiments and analysed and interpreted the data. Z.A. performed the RT-qPCR for Figures 1C and S1F. S.A. helped with the analysis of the DAP-seq dataset for Figures 3C and 4C. S.C and J.C.W.L wrote the article, with S.C writing the first draft‥

